# Cross-talk between RNA secondary and three-dimensional structure prediction: a comprehensive study

**DOI:** 10.1101/2025.06.23.661015

**Authors:** Deyin Wang, Yangwei Jiang, Linli He, Linxi Zhang, Ruhong Zhou, Dong Zhang

## Abstract

In recent years, various computational methods have been developed to predict the three-dimensional (3D) structures of RNAs. Due to its hierarchical folding property, RNA secondary (2D) structure is usually used as input for 3D structure prediction to improve accuracy and efficiency. However, the extent to which the accuracy of input 2D structure affects the performance of 3D structure prediction remains to be further investigated. Additionally, whether and how the input base-pairing interactions are modified during the 3D structure modeling process is another question worth exploring. To address these issues, here we comprehensively benchmark five representative 3D structure prediction models on extensive datasets, using 2D structures of varied accuracies as input. Our results indicate that there is a pervasive cross-talk between RNA 2D and 3D structure predictions, where the performance dependence of 3D structure prediction on the accuracy of input 2D structure is closely associated with the 3D model’s ability to modify the input base-pairing interactions during structure modeling. Furthermore, we also observed that RNA 3D structure prediction performance is more sensitive to the occurrence of false positive base pairs in the input 2D structure than to true positive base pairs, suggesting a worthy direction to further improve the model performance.

**Author summary:** Three-dimensional (3D) structural modeling of RNAs with large sizes and complex topologies remains challenging despite the availability of (predicted) 2D structures as constraints. Exploring the potential cross-talk between 2D and 3D structure predictions is a worthy direction to improve the performance of RNA structure modeling. In this study, we found that all tested popular RNA 3D structure prediction models were able to modify the original base pairing interactions contained in the input 2D structure during the 3D modeling process. Especially, all of these models presented increased F1-score for the optimal combination of RNA 2D and 3D structure prediction models. The results suggest that a worthwhile direction to further improve the performance of RNA 3D structure prediction is to minimize the incidence of incorrectly predicted base pairing interactions during modeling process without compromising or even improving the presence of correct interactions in the input 2D structure.

## Introduction

Ribonucleic acids (RNAs) are important functional macromolecules in living organisms. The particular three-dimensional (3D) structure adopted by RNA molecules is crucial for their biological functions. However, determining the actual 3D structure of RNA often requires expensive and time-consuming experimental techniques. Therefore, computer programs for RNA 3D structure prediction have emerged in the past three decades [1,2]. Current RNA 3D structure prediction methods primarily fall into three categories: template-based methods, *de novo* methods, and the most recent deep learning-based methods. Template-based methods rely on known structures/fragments as templates to predict RNA 3D structures, including 3dRNA [3], ModeRNA [4], RNAComposer [5], Vfold3D [6], FARFAR2 [7], and more [8–13]. They search for templates with similar sequence or structural characteristics to model the 3D structure of the target molecule. On the other hand, *de novo* methods can predict the 3D structures of RNAs from scratch based on sequence information, with or without secondary (2D) structure as constraints, such as HiRE-RNA [14], IsRNA [15–17], iFoldRNA [18], SimRNA [19], and more [20–25]. They utilize physical and chemical principles to sample the conformational space and predict the most stable/probable conformation of the target molecule. Finally, inspired by the success of deep learning-based protein 3D structure prediction in the past few years [26–29], many deep learning-based methods have been developed to facilitate RNA 3D structure prediction, such as DRfold [30], RhoFold+ [31], RoseTTAFoldNA [32], trRosettaRNA [33] and more [34–36]. Nevertheless, compared with protein structure prediction, these deep learning-based methods generally perform much worse for RNAs, indicating that RNA 3D structure prediction remains challenging [37].

In parallel, a collective and blind experiment called RNA-Puzzles was launched to assess the leading edge of RNA structure prediction technique [38–42]. Their results emphasized that computational methods for RNA structure prediction can already provide useful structural information for biological problems, but the poor prediction of non-Watson-Crick interactions suggests that the algorithms need further improvement. Meanwhile, in 2022, twelve experimental RNA targets were first introduced into CASP15 [43]. Two different assessments independently ranked four traditional methods (template-based or *de novo* methods) as top predictors, while deep learning-based approaches performed significantly worse than these top-ranked groups. Particularly, for synthetic RNA targets in CASP15 that lack homologous RNA sequences and structures akin to existing RNAs, their precise 3D structure modeling necessitates significant interventions from human experts [44–47]. Therefore, it is crucial to conduct more comprehensive benchmarking to identify the critical factors for predicting RNA 3D structures. Tentative benchmarking of RNA 3D structure prediction tools have been carried out recently [48–50], but these studies were limited by a narrow selection of RNA datasets or focusing exclusively on deep learning methods. We anticipate that insights gained from more comprehensive benchmark tests, combined with continued advances in computational biology, available datasets, and deep learning algorithms, may address the challenges of accurate RNA 3D structure prediction in the future.

In general, since RNA folding is hierarchical [51,52], its 2D structure is usually used as a constraint in 3D structure modeling to improve prediction efficiency and accuracy. So far, numerous models have been proposed to obtain the RNA 2D structure from its sequence, such as RNAfold [53], RNAStructure [54], CONTRAfold [55], Mfold [56–58], HotKnots [59], MXfold2 [60], NUPACK [61], SPOT-RNA [62], and more [63–69]. However, the 2D structure prediction accuracy of existing models is usually limited [70,71]. For example, on datasets containing diverse RNA sequences, the average F1-score of almost all tested models is less than 0.8 [60,62]. Thus, an extensive benchmark is required to examine how these predicted 2D structure inputs affect the prediction accuracy of current 3D structure methods. Furthermore, whether and how the input 2D structure is changed during the 3D structure modeling process is another question worth exploring.

Here, we comprehensively benchmark the performance of several representative RNA 3D structure prediction methods based on an extensive array of datasets, including RNAs considered in CASP15, CASP16, RNA-puzzles, and a custom dataset composing of 31 curated RNAs. Our goals include: (1) comparing the prediction performances of typical RNA 3D methods, including template-based, *de novo*, and deep learning-based methods, using the native 2D structure as input; (2) exploring the impact of 2D structure prediction accuracy on RNA 3D structure modeling, where 2D structures predicted by different models were used as inputs; (3) providing useful guidelines to facilitate the optimal combination of different 2D and 3D structure prediction methods; and (4) examining whether and how the raw 2D structure input is altered during the 3D structure modeling process. In addition to evaluating the prediction accuracy of different methods, we anticipate that the study of the cross-talk between RNA 2D and 3D structure predictions can reveal potential factors that affect the accuracy of RNA structure prediction, thereby paving the way for improving the accuracy and reliability of future computational models.

## Results

### Benchmark results for RNA 3D structure prediction based on native 2D structure

To explore the prediction upper bounds of the five selected RNA 3D models (see Table 1), we first performed benchmark tests on three different datasets, including “Custom”, “RNA Puzzles”, and “CASP RNA” (see Methods and Materials and Supplementary Information (SI) Tables S1-S3 for details), using the native 2D structure as input. As shown in Fig 1, for the Custom dataset, the template-based model RNAComposer provides the best predictions in terms of the RMSD metric, as it gives the lowest median RMSD = 13.7 Å (see Fig 1A) and a relatively high median INF_ALL = 0.78 (see Fig 1B). However, in terms of the TMscore metric, IsRNA2 and FARFAR2 show a significant leading performance with median TMscore = 0.27 (see Fig 1C). In addition, DRfold provides the best predictions in terms of lDDT (median lDDT = 0.64, see Fig 1D). These results re-emphasize the complexity of evaluating RNA 3D structure predictions and the necessity of using multiple metrics. For the RNA Puzzles dataset, the *de novo* IsRNA2 method provides the best predictions for almost all metrics among the five selected 3D models (median RMSD = 8.6 Å, median INF_ALL = 0.83, median TMscore = 0.32, and median lDDT = 0.64). While for the CASP RNA dataset, RNAComposer ranks top among the five selected models (median RMSD = 18.1 Å, median INF_ALL = 0.84, median TMscore = 0.29, and median lDDT = 0.61). Furthermore, we also considered the predictions from AlphaFold3 and trRosettaRNA as a reference, which show leading performance on almost all three test datasets (see Fig 1). The full list of RMSD for all tested methods is available in Table S4. However, due to the possible data overlap between the training set of AlphaFold3 and trRosettaRNA and the test datasets constructed here, we will not discuss the results of AlphaFold3 and trRosettaRNA in depth in subsequent study. Overall, although the prediction performance of the five representative 3D models for different RNA targets varied when using the native 2D structures as input, the predictions of traditional methods such as RNAComposer and IsRNA2 methods are generally promising, which is consistent with the observations in the recent CASP15 competition [43].

**Table 1.** Overview of RNA 3D structure prediction models tested in this study.

| **Model** | **Category** | **Access** | **Inputs** |
| --- | --- | --- | --- |
| **DRfold [30]** | Deep Learning-based | Local | Seq+2D |
| **RNAComposer [5,46]** | Template-based | Webserver | Seq+2D |
| **FARFAR2 [7]** | Template-based | Local | Seq+2D |
| **IsRNA2 [17]** | *de novo* | Local | Seq+2D |
| **SimRNA [19]** | *de novo* | Local | Seq+2D |

**Fig 1.**
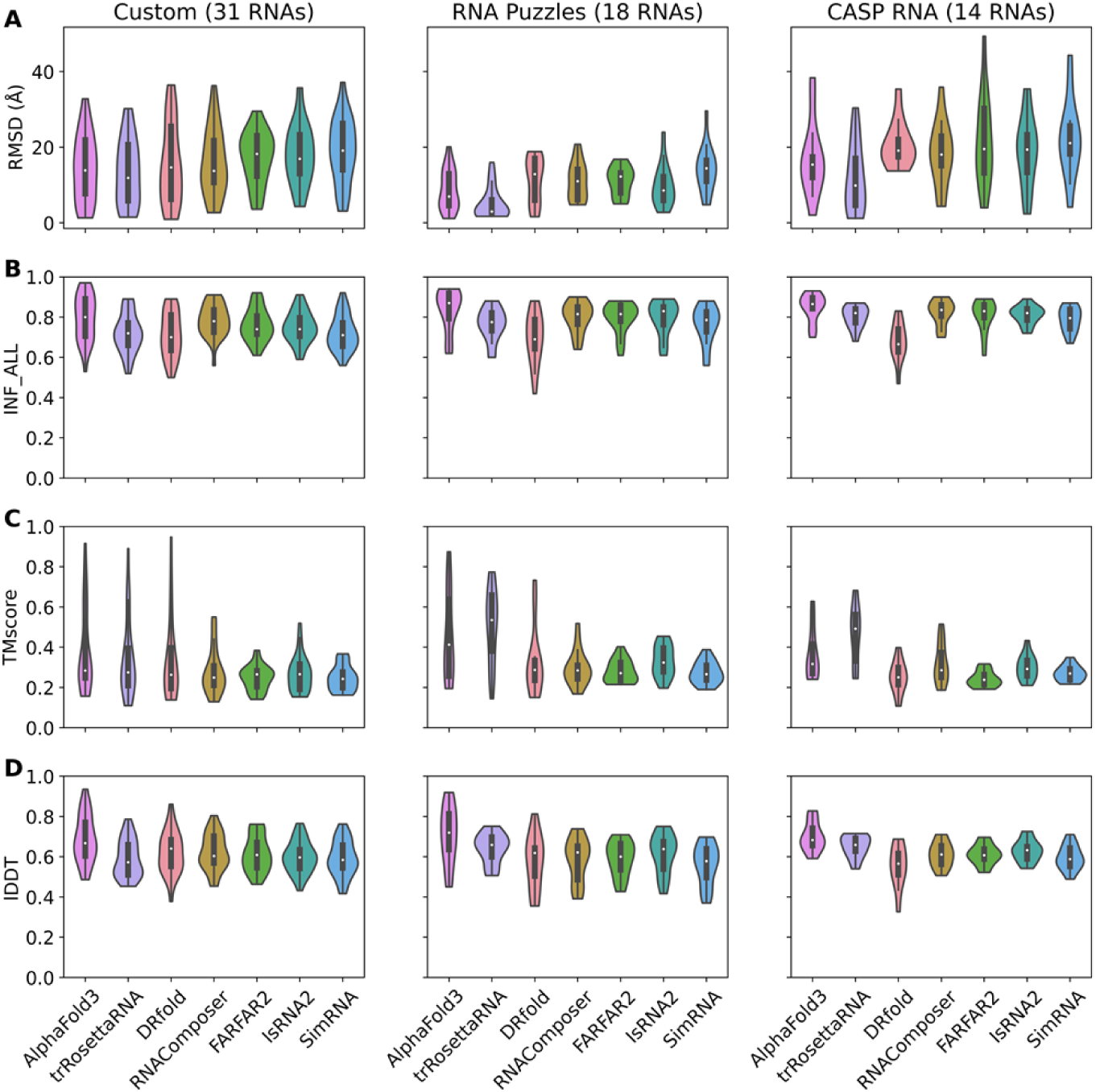
Benchmark results of RNA 3D structure prediction using the native 2D structure as input. Violin plots of (A) RMSD, (B) INF_ALL, (C) TMscore, and (D) lDDT predicted by five selected 3D structure models. Three different test datasets were benchmarked: a custom dataset collected from PDB (first column), the real challenge RNA Puzzles (second column) and CASP RNA (third column). The number of RNAs contained in each dataset is shown in parentheses.

### Benchmark results based on 2D structures predicted by different models

Now we consider a more general RNA 3D structure prediction scheme: using a pre-generated 2D structure as input to predict the possible 3D conformation of the query sequence. For a comprehensive benchmarking, we tested six popular 2D structure prediction tools (see Table 2 for details) to generate the input 2D structure. To facilitate subsequent analysis, we first investigated the 2D structure prediction accuracy of different 2D models. Since the RNA structure prediction accuracy (both 2D and 3D) usually depends on its structural topology, here we merged the above three test datasets into a combined dataset and then reclassified them into three categories based on their native structural information for in-depth analysis: stem-loops, multi-way junctions, and pseudoknots containing tertiary interactions (see Fig S1). As listed in Table 3, all six tested 2D tools have limited prediction accuracy, indicating that accurate RNA 2D structure prediction remains challenging. For instance, the best mean F1-scores are 0.819 (by MXfold2), 0.857 (by MXfold2), and 0.810 (by CONTRAfold) for stem-loops, multi-way junctions, and pseudoknots, respectively. Interestingly, when we considered 2D structures derived from AlphaFold3 RNA 3D structure predictions, we observed a significant improvement in 2D structure accuracy for all three structural categories (with mean F1-score > 0.91). This suggests a potential avenue to address the long-standing bottleneck of RNA 2D structure prediction, e.g., the accuracy of 2D structure prediction can be improved by taking tertiary structures into account. Regardless, due to their relatively high quality, AlphaFold3-derived 2D structures were also used as another baseline input (in addition to the native 2D structures) to extend our benchmark results in subsequent studies. As shown in Fig S2, the combination of different 2D and 3D structure prediction models leads to varied RNA 3D structure prediction performances. Systematic analysis of these combination results can provide useful guidance for constructing a well-tuned RNA 3D structure modeling pipeline, such as using AlphaFold3-derived 2D structures as input for IsRNA2 RNA 3D structure prediction is optimal, while using native2D structures is suboptimal.

**Table 2.** Summary of RNA 2D structure prediction models tested in this study.

| **Model** | **Category** | **Access** |
| --- | --- | --- |
| **RNAfold [53]** | Thermodynamics | Local |
| **NUPACK [61]** | Thermodynamics | Local |
| **Mfold [56–58]** | Thermodynamics | Local |
| **RNAstructure [54]** | Thermodynamics | Local |
| **CONTRAFold [55]** | Deep Learning | Local |
| **MXfold2 [60]** | Deep Learning | Local |

**Table 3.** Summary of F1-scores for predictions from different 2D models. Mean values for RNAs with different topologies were displayed. The number of RNAs involved in each category is given in parentheses.

|  | **Stem-loop**<br>**(22)** | **Multi-way junction**<br>**(16)** | **Pseudoknot**<br>**(25)** |
| --- | --- | --- | --- |
| **RNAfold** | 0.694 | 0.773 | 0.737 |
| **NUPACK** | 0.667 | 0.580 | 0.710 |
| **Mfold** | 0.702 | 0.829 | 0.691 |
| **RNAstructure** | 0.731 | 0.783 | 0.757 |
| **CONTRAFold** | 0.704 | 0.701 | 0.810 |
| **MXfold2** | 0.819 | 0.857 | 0.765 |
| **AlphaFold3** | 0.914 | 0.961 | 0.970 |

As shown in Fig 2, when using predicted 2D structures as input, the five tested 3D models performed similarly on stem-loops with median RMSD = 19.5-21.1 Å (see Fig 2A), median INF_ALL = 0.66-0.68 (see Fig 2B), and median TMscore = 0.22-0.24 (see Fig 2C). While for multi-way junctions and pseudoknot RNAs, IsRNA2 and DRfold performed best, with median RMSD = 17.2 and 15.9 Å (see Fig 2A), respectively. We also noticed that 3D structure predictions on stem-loops are slightly worse than those on multi-way junctions and pseudoknots, i.e., the median RMSD value for stem-loops is relatively large, even though the latter two categories are generally more structurally complex than the former. The relatively high proportion of unpaired nucleotides in the tested stem-loop RNAs (see Fig S1C) may be responsible for this phenomenon.

**Fig 2.**
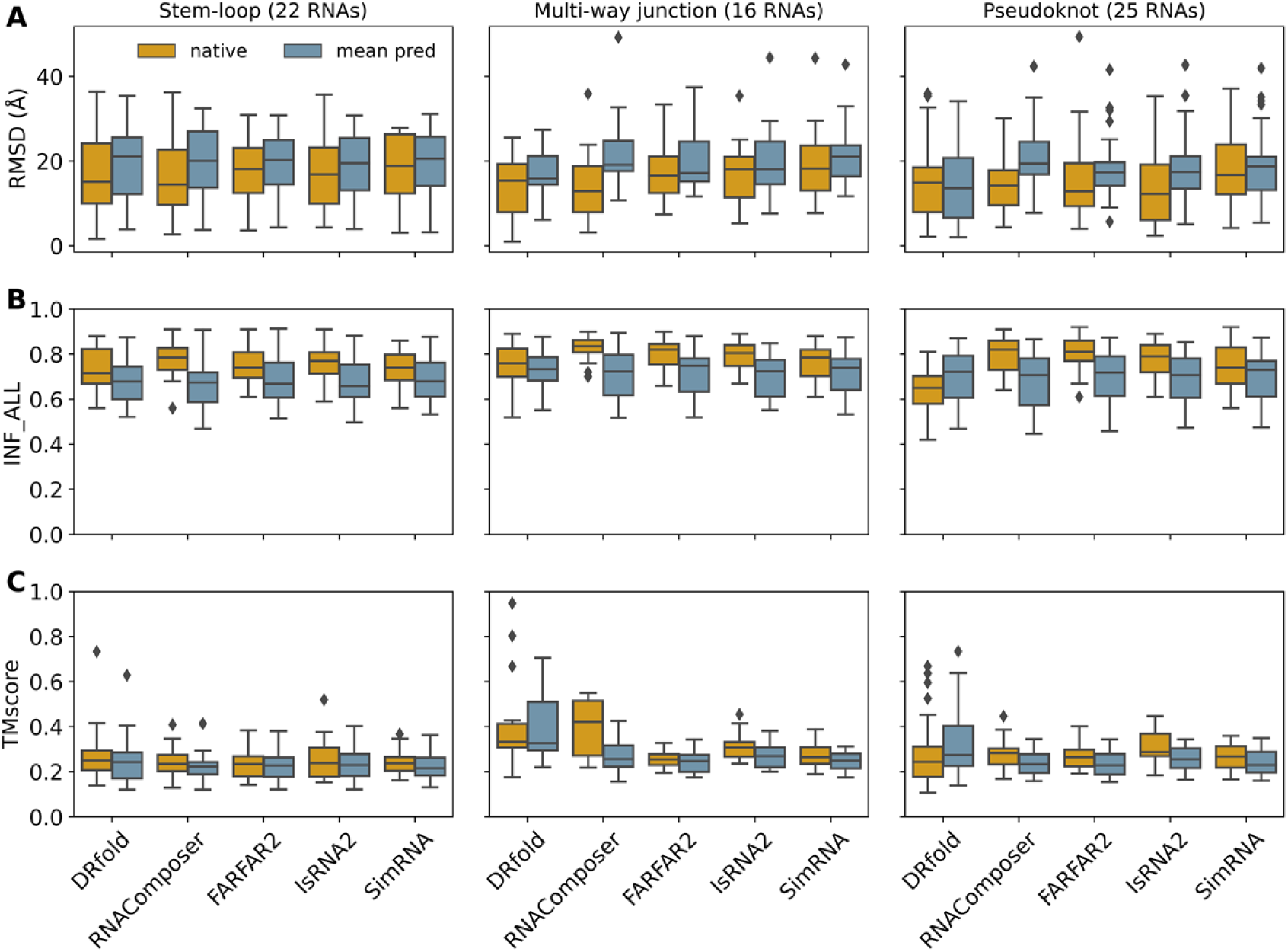
Summary of RNA 3D structure prediction performance using different 2D structures as input. Box plots of (A) RMSD, (B) INF_ALL, and (C) TMscore values for the predictions of five tested 3D models on different topological categories: stem-loop RNAs (first column), RNAs containing multi-way junctions (second column), and RNA pseudoknots containing tertiary interactions (third column). Predictions using native 2D structures as input are denoted as “native”, and “pred” represents the values of 3D structure predictions using different 2D structures generated by the six selected 2D tools as input.

As shown in Fig 2, for almost all five tested 3D models, RNA 3D structure predictions using the predicted 2D structures as input generally performed worse than predictions based on the native 2D structures, i.e., the former usually have relatively higher median RMSDs, relatively lower median INF_ALL and TMscore values. Specifically, for RNAComposer and IsRNA2, the difference in prediction performance between the two cases (using the native *vs.* predicted 2D structure as input) is significant on all three structural categories; while for SimRNA on stem-loop RNAs and DRfold on multi-way junctions, the difference is less pronounced (see Fig 2). Unexpectedly, for DRfold on pseudoknot RNAs containing tertiary interactions, we found that predictions using predicted 2D structures as input outperformed those based on the native 2D structures. This counterintuitive result may partly come from the fact that DRfold used predicted 2D structures (by RNAfold and PETfold) rather than the native 2D structure to prepare features during model training. In addition to the average performance over 3D predictions using all predicted 2D structures by the six selected 2D tools as input, we also analyzed 3D structure predictions based only on the optimal predicted 2D structure or the AlphaFold3-derived 2D structure, and the results are shown Fig S3. Overall, these results preliminarily demonstrated that different 3D models have varied dependencies on the accuracy of the input 2D structure.

### Dependence of 3D structure prediction performance on 2D structure accuracy

In general, the accuracy of 2D structures generated by currently available 2D tools is limited, and the accuracy may vary for different RNA sequences (see Table 3). Therefore, how these imperfect 2D structures (with F1-score < 1.0) as input affect the prediction performance of RNA 3D structure prediction methods is a question worthy of further exploration. To address this, we investigated the relationship between the RMSD/INF_ALL values of predicted 3D structures and the F1-score values of input 2D structures in the Combined dataset. As shown in Fig 3, for all five tested 3D methods, when the F1-score increases from 0 to 1.0, the overall trend of the RNA 3D structure prediction accuracy is on the rise (see Fig 3A), where RMSDs tend to decrease gradually and INF_ALL values tend to increase gradually (see Fig 3B). Some local fluctuations on the profiles of RMSD/INF_ALL may be caused by the uneven distribution of different topological categories in particular bins of F1-score values (see Fig S4). In terms of individual test datasets, we observed similar variation trends for RMSD and INF_ALL in the Custom and RNA Puzzles datasets (see Fig S5). These results indicated that more accurate 2D structure as input can generally improve the performance of RNA 3D structure prediction. Furthermore, we found that DRfold significantly outperformed the other four 3D models in terms of RMSD and INF_ALL in the interval of F1-score = 0.55-0.75, suggesting its unique dependence on the accuracy of input 2D structure, which will be further discussed below.

**Fig 3.**
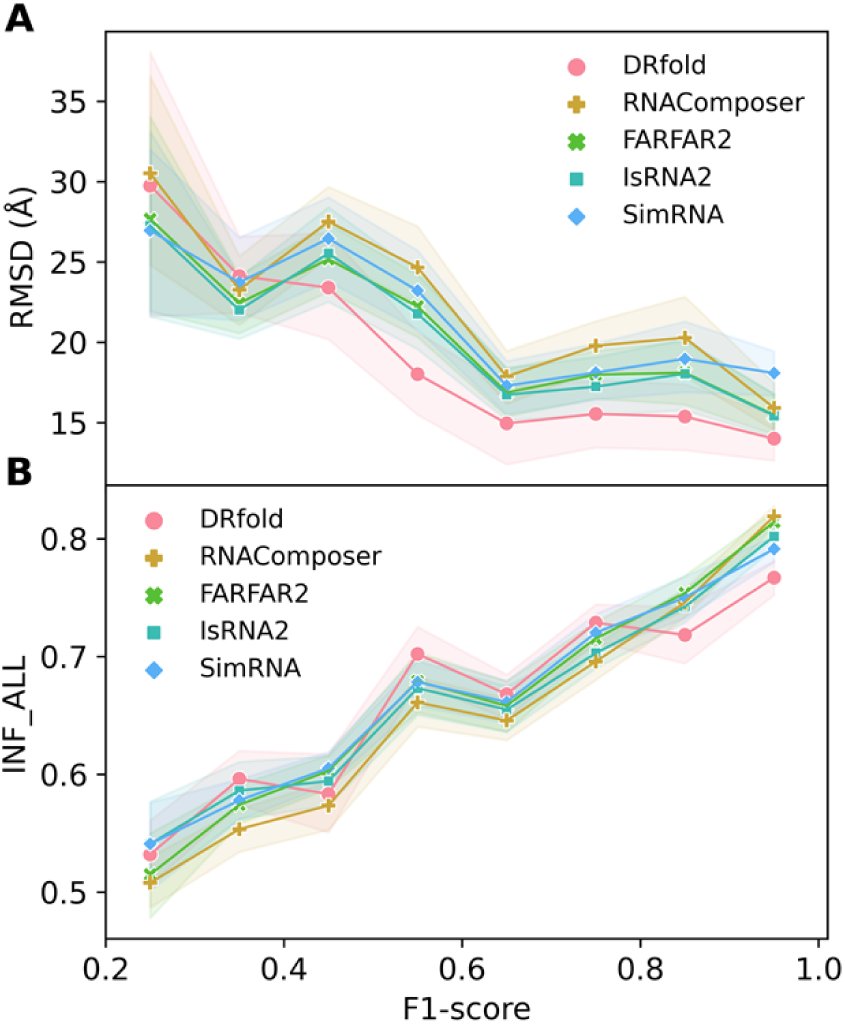
Quality of 3D structure predictions depends heavily on the accuracy of input 2D structures. (A) RMSD and (B) INF_ALL values of RNA 3D structures predicted by the five selected models as functions of the F1-score values of input 2D structures. The F1-score values were grouped by a bin size of 0.1. Symbols represent the mean metrics of a particular bin and shaded area indicates the 95% confidence interval. The analysis considered all 2D structures predicted by the selected six 2D tools, AlphaFold3-derived 2D structures, and the native 2D structures.

To further investigate to what extent the accuracy of the input 2D structure affects the performance of individual tested RNA 3D structure prediction models, we classified the predicted 2D structures into three categories according to their F1-score: low accuracy (F1-score < 0.5), medium accuracy (0.5 ≤ F1-score < 0.9), and high accuracy (F1-score ≥ 0.9). As shown in Fig S6, for 2D structures with low accuracy as input, the accuracy of 3D structure predictions for the majority of RNAs is generally poor. Among the five selected 3D models, the best median RMSD is 19.8 Å (obtained by IsRNA2), the highest median INF_ALL is 0.57 (obtained by SimRNA), and the highest median TMscore is 0.18 (obtained by SimRNA). In contrast, when using high-accuracy 2D structures as input, the quality of 3D structure predictions improves apparently, with the best median RMSD dropping to 13.2 Å (obtained byDRfold), the best INF_ALL increasing substantially to 0.83 (obtained by RNAComposer), and the best median TMscore also increasing to 0.30 (obtained by IsRNA2 and DRfold). Interestingly, when the accuracy of input 2D structure is moderate, DRfold could still provide acceptable 3D structure predictions for some RNAs (with median RMSD/INF_ALL/TMscore = 16.0 Å/0.72/0.29), and its performance is clearly better than that of the other four 3D models. Taken together, these results indicated that when the input 2D structure is highly confident (very close to the native one), the 3D structure prediction by IsRNA2 is generally more reliable, which is also consistent with the results shown in Fig 1. When the input 2D structure is only of medium accuracy, the 3D structure predicted by DRfold is strongly recommended, which is in good agreement with results displayed in Figs 2 and 3; while for the input 2D structure with low accuracy, the 3D structure predictions by IsRNA2, DRfold, and SimRNA seem relatively more promising.

There are two different types of base-pairing interactions in the predicted 2D structures: true positive base pairs that are correctly predicted and false positive base pairs that have no correspondence in the native structure. To deepen the understanding of the relationship between the accuracy of 3D structure prediction and the accuracy of input 2D structure, we then analyzed the relationship between the prediction quality (characterized by RMSD and INF_ALL) and α_*TP*_ (see Eq. 1) or α_*FP*_ (see Eq. 2) for each tested 3D model. As shown in Fig 4, for all five 3D methods, the INF_ALL values are nearly positively correlated with α_*TP*_ and negatively correlated with α_*FP*_, which again declared that the accuracy of base pairs in the input 2D structure has a significant influence on the final accuracy of the predicted 3D structure. Expectedly, the positive correlation of INF_ALL with α_*TP*_ and the negative correlation of RMSD with α_*TP*_ (see Fig S7) indicated that the more correctly the base-pairing interactions in the input 2D structure were predicted, the higher the likelihood of achieving high-accuracy RNA 3D structure prediction. Conversely, the negative correlation of INF_ALL with α_*FP*_and the positive correlation of RMSD with α_*FP*_ (see Fig S7) suggested that incorrectly predicted base-pairing interactions in the input 2D structure often lead to lower-accuracy 3D structure models. In general, those false positive base pairs introduce noise and mislead 3D structure modeling algorithms by suggesting incorrect folding patterns or interaction networks, especially for the template-based methods. Additionally, we observed that predictions by RNAComposer, FARFAR2, and IsRNA exhibited higher dependence of INF_ALL on α_*TP*_ and α_*FP*_ (Pearson correlation coefficient *R* ≥ 0.66), while predictions of DRfold showed distinctly reduced dependence (*R* = 0.46), as illustrated by the fitting lines in Fig 4A; see also Fig S7 for the dependence of RMSDs on α_*TP*_ and α_*FP*_. These results suggested that the more sophisticated and accurate deep learning-based geometrical potentials [30] for 3D structure prediction do not rely heavily on the accuracy of input 2D structure compared to traditional methods. Furthermore, although the number of false positive base pairs exceeds the number of native base pairs (α_*FP*_ > 1.0) in some cases, the INF_ALL value did not display an abnormal drop (see Fig 4B), indicating the robustness of tested 3D models in structure prediction.

**Fig 4.**
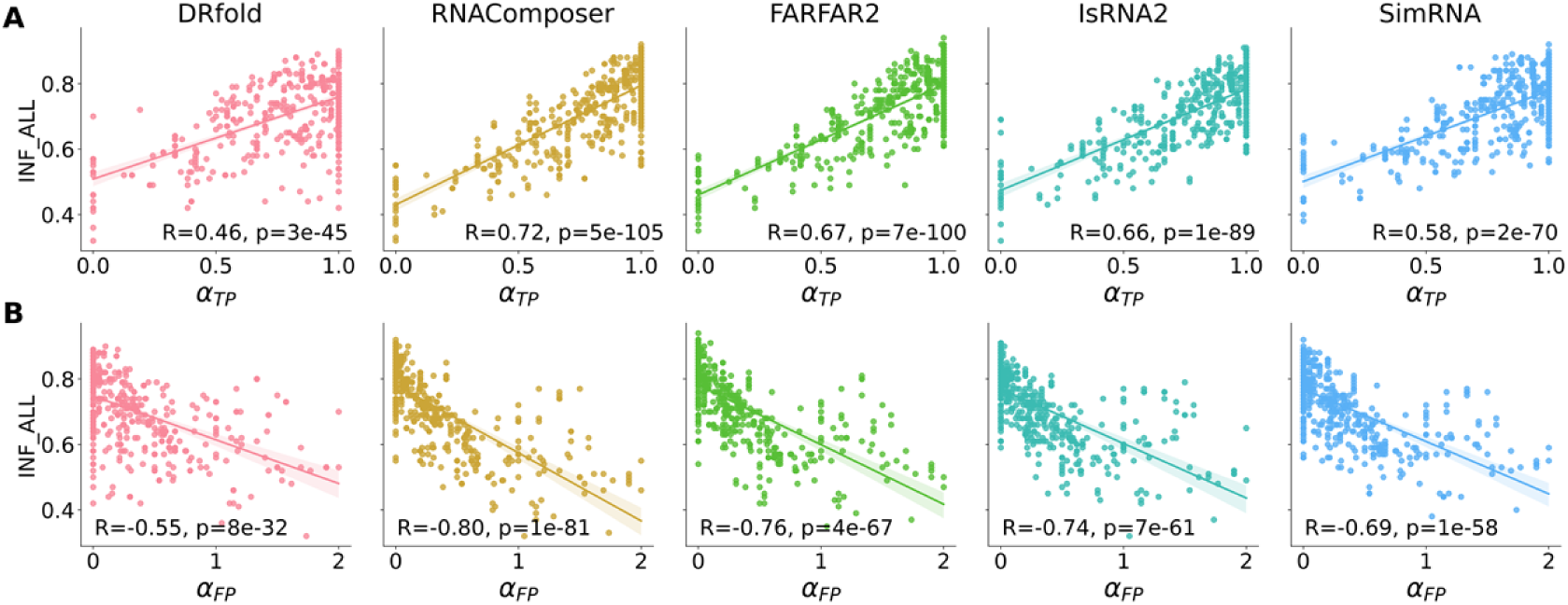
Different base pair compositions in the input 2D structure have distinct influences on the quality of 3D structure predictions. Scatter plots shown the relationship between the (A) proportion of true positive (α_*TP*_) base pairs and (B) proportion of false positive (α_*FP*_) base pairs in the input 2D structure and the INF_ALL values of 3D structures predicted by different 3D models. The line in each plane represents the linear fitting result, and the corresponding Pearson correlation coefficient (*R*) and *p*-value (*p*) are presented in the bottom. From left to right: DRfold, RNAComposer, FARFAR2, IsRNA2, and SimRNA.

### Modification of base-pairing interactions during 3D structure modeling

In Fig 4, we noticed that even if the true positive base pairs in the input 2D structure are zero (α_*TP*_ = 0), the INF_ALL values of predictions from all five tested 3D models are non-zero, and some predictions even have INF_ALL > 0.5. This indicated that all these 3D models are able to (partially) modify the input base-pairing interactions during the 3D structure modeling process, e.g., identifying and forming correct base pairs. A similar phenomenon has also been reported in other study [33]. Two illustrative examples were presented in Fig 5 to further demonstrate this observation. The predicted 2D structures of these two RNAs as input deviate markedly from their corresponding native 2D structures (F1 = 0 and 0.19, respectively). However, SimRNA (see Fig 5A) and DRfold (see Fig 5B) still generated high-quality 3D structure predictions for these two RNAs with RMSD ≤ 3.6 Å and INF_ALL ≥ 0.64, respectively. One possible reason for this capability is that many important functional motifs in RNAs are usually conserved, and RNA 3D structure prediction models might be biased towards these motifs (either through data training or existed templates or knowledge-based potentials). Therefore, even if the input 2D structure is far from the native 2D structure, some key tertiary interactions can still be recovered during RNA 3D structure modeling. In addition, we also observed cases where base-pairing interactions deteriorated during 3D structure modeling; see SI Text and Fig S8 for more details.

**Fig 5.**
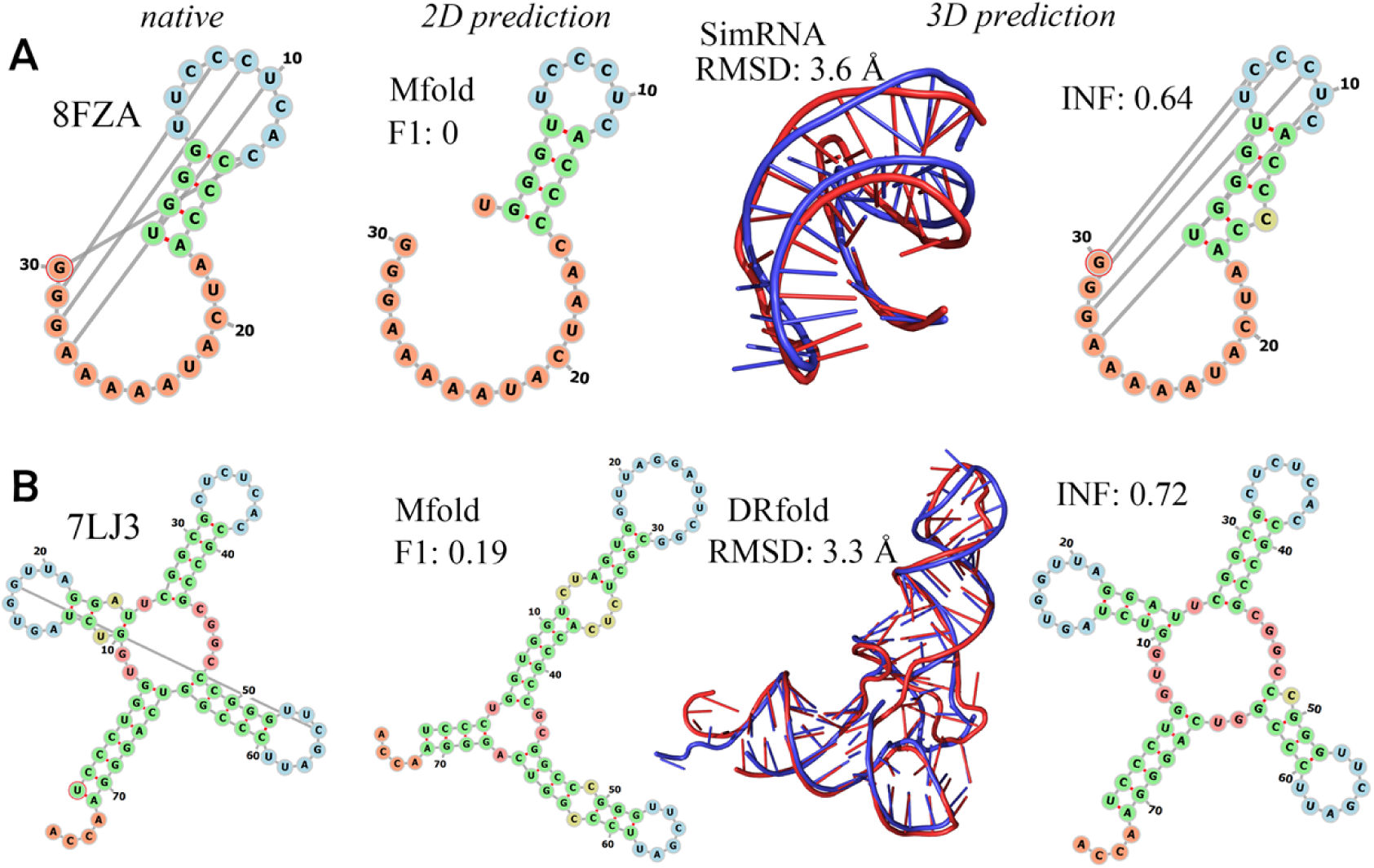
Illustrative examples of modifications of base-pairing interactions during 3D structure modeling when using an incorrected 2D structure as input. (A) The Class I type III preQ1 riboswitch from *E. coli* [80] (PDB id: 8FZA) and its 3D structure prediction by SimRNA. (B) The structure of human transfer RNA visualized in the cytomegalovirus [81] (PDB id: 7LJ3) and its 3D structure prediction by DRfold. Columns from left to right: the native 2D structure extracted from experimental structure, predicted 2D structure by Mfold as input, predicted 3D structure (in red) aligned with experimental structure (in blue), and modified 2D structure derived from the predicted 3D structure. RMSD and INF_ALL values between the predicted and native structures are also shown.

To gain a preliminary understanding of the ability of various 3D models to modify the input base-pairing interactions during RNA 3D structure modeling, we analyzed the changes in the interaction network of predicted 3D structures using 2D structures of different accuracy as input. Fig 6 showed the changes in F1-score (ΔF1), proportion of true positive base pairs (Δα_*TP*_), and false positive base pairs (Δα_*FP*_) relative to the input 2D structure during 3D structure modeling for all five selected 3D models. To understand the better performance of deep learning-based RNA 3D structure prediction methods, we also showed the results of trRosettaRNA tested on our datasets in Fig 6. Overall, for DRfold, many predictions improved their base-pairing interactions relative to the input 2D structure (ΔF1 > 0, see Fig 6A), especially for inputs with F1 < 0.8, which was caused by the introduction of more true positive base pairs (Δα_*TP*_ > 0, see Fig 6B) and/or the elimination of false positive base pairs (Δα_*FP*_ < 0, see Fig 6C) during structure modeling. We also observed similar results for some predictions of IsRNA2 and SimRNA when using low-accuracy 2D structures (F1 < 0.3, see Fig 6A) as input. Furthermore, when using high-accuracy 2D structures as input, we also found that some predictions from IsRNA2, SimRNA and RNAComposer had deteriorated base-pairing interactions (ΔF1 < 0), i.e., a decrease in true positive base pairs (Δα_*TP*_ < 0) and an increase in false positive base pairs (Δα_*FP*_ > 0); see Fig 6. These results indicated that modification of input base-pairing interactions during structure prediction is prevalent in the tested 3D models, and that this modification capability varies between different 3D models.

**Fig 6.**
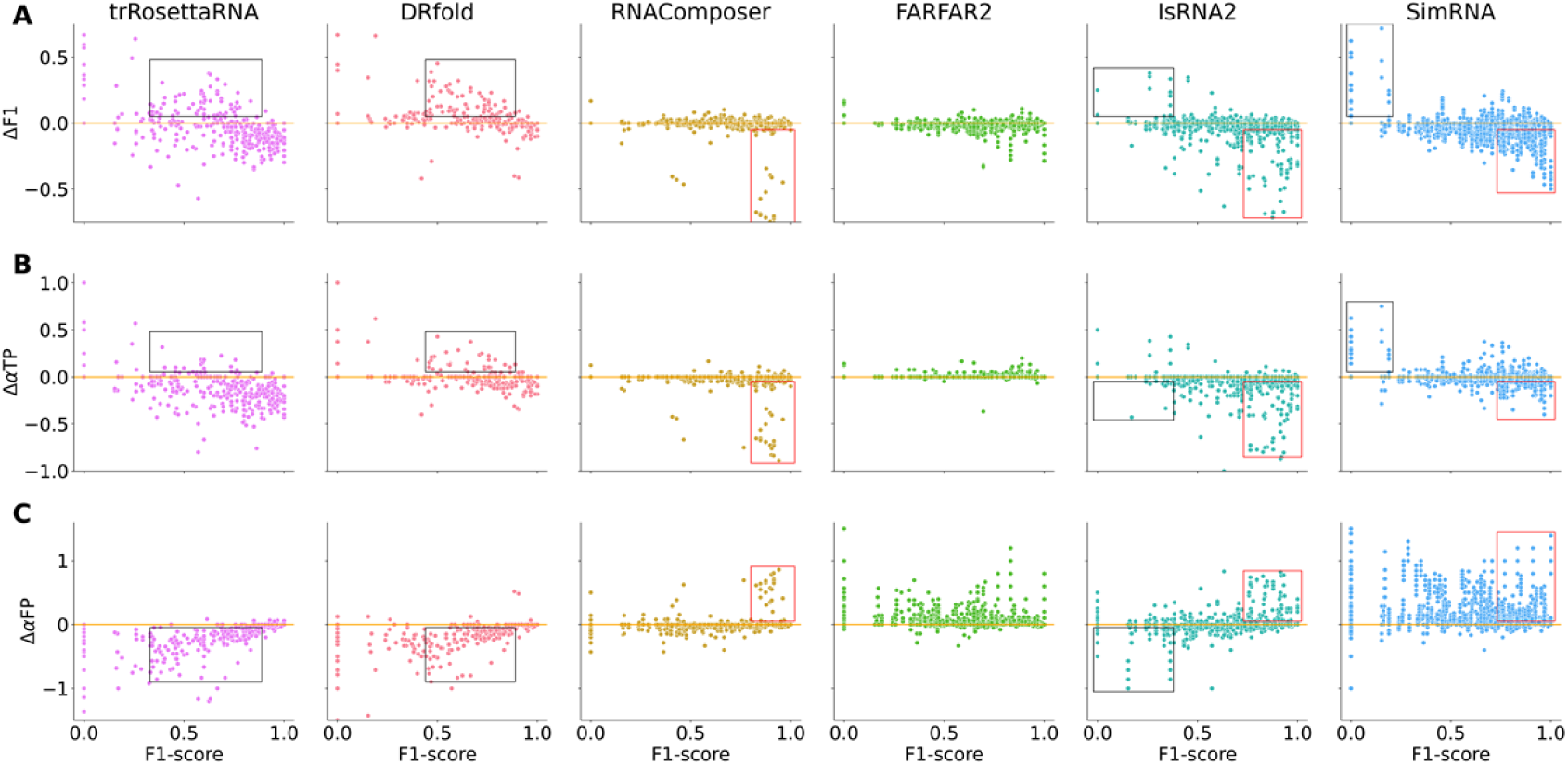
Different 3D models vary in their ability to modify input base-pairing interactions during structure prediction. Scatter plots of the changes in (A) F1-score (ΔF1), (B) proportion of true positive (Δα_*TP*_), and (C) proportion of false positive (Δ α_*FP*_) base pairs during 3D structure modeling *vs.* F1-score of the input 2D structure. Each point represents a 3D structure prediction using a 2D structure generated by one of the six selected 2D tools as input. Predictions with notably modified base-pairing interactions are marked with rectangular boxes. From left to right: trRosettaRNA, DRfold, RNAComposer, FARFAR2, IsRNA2, and SimRNA.

Finally, we attempted to establish a link between the 3D structure prediction performance of each tested 3D model and its ability to modify input base-pairing interactions during the RNA 3D structure modeling process, namely cross-talk between RNA 2D and 3D structure predictions. To this end, we compared the changes in base-pairing interactions during 3D structure predictions when using different 2D structures as input, including the optimal 2D structure that produced the best prediction (in terms of RMSD) among the six selected 2D tools, the AlphaFold3-derived high-accuracy 2D structure, and the native 2D structure. As shown in Fig 7, for the template-based RNAComposer, the ability to modify the input base-pairing interactions was negligible in most predictions (median ΔF1 ∼ 0.01), regardless of whether the AlphaFold3-derived or native 2D structures were used as input. This point is consistent with the above observation that the performance of RNAComposer is heavily dependent on the accuracy of input 2D structure. For example, RNAComposer performs well when using the native 2D structure as input (see Fig 1). For the *de novo* IsRNA2 model, many predictions can eliminate some incorrect base-pairing interactions when using the optimal 2D structures as input (median Δα_*FP*_ = ―0.07, see Fig 7B, and ΔF1 = 0.02, see Fig 7A). Correspondingly, we observed that the accuracy of input 2D structure plays a key role in the 3D structure prediction performance of IsRNA2. For instance, IsRNA2 performs top among the five selected 3D models in the benchmark based on the native 2D structure (see Fig 1); while when using the predicted 2D structures as input, its performance is comparable to but slightly worse than the best model (see Figs 2 and S3). For the deep learning-based methods DRfold and trRosettaRNA, we observed clearly distinct results when using different 2D structures as input. Specifically, when the optimal predicted 2D structure (usually with medium accuracy) was used as input, predictions of DRfold and trRosettaRNA can improve the quality of base-pairing interactions in many cases (median ΔF1 = 0.07 and 0.04, see Fig 7A), which was mainly achieved by eliminating some erroneous base pairs during structure modeling (median Δα_*FP*_ = ―0.22 and -0.28, see Fig 7C). This signature distinguishes deep learning-based methods from the other four tested 3D models, which may explain its unique dependence on the accuracy of input 2D structure in the interval of F1-score = 0.55-0.75 (see Fig 3). However, when using the AlphaFold3-derived or native 2D structures (with high accuracy) as input, 3D structure predictions by DRfold generally worsened the input base-pairing interactions (median ΔF1 < 0, see Fig 7A) through excluding some true positive base pairs (median Δα_*TP*_ < 0, see Fig 7B) and/or introducing some false positive base pairs (median Δα_*FP*_ > 0, see Fig 7C). This phenomenon may account for its relatively mediocre 3D structure prediction performance in the test using the native 2D structure as input (see Fig 1), as well as the better performance using predicted 2D structures as input than those using the native 2D structure as input for pseudoknot RNAs (see Fig 2). Furthermore, we also compared the F1-scores between the predicted 3D structures and their corresponding 2D inputs and observed consistent results. Namely, all five tested 3D models could present predictions where the F1-score of generated 3D structure is greater than the corresponding 2D input (in a fraction of 0.24-0.52; see Fig S9). Meanwhile, predictions of RNAComposer had the most cases (fraction > 0.5) where the F1-score did not change, while DRfold’s predictions had the least cases. Overall, these results declared that the cross-talk between RNA 2D and 3D structure predictions is pervasive and that the dependence of 3D structure prediction performance on the accuracy of input 2D structure is closely associated to the model’s ability to modify the input base-pairing interactions.

**Fig 7.**
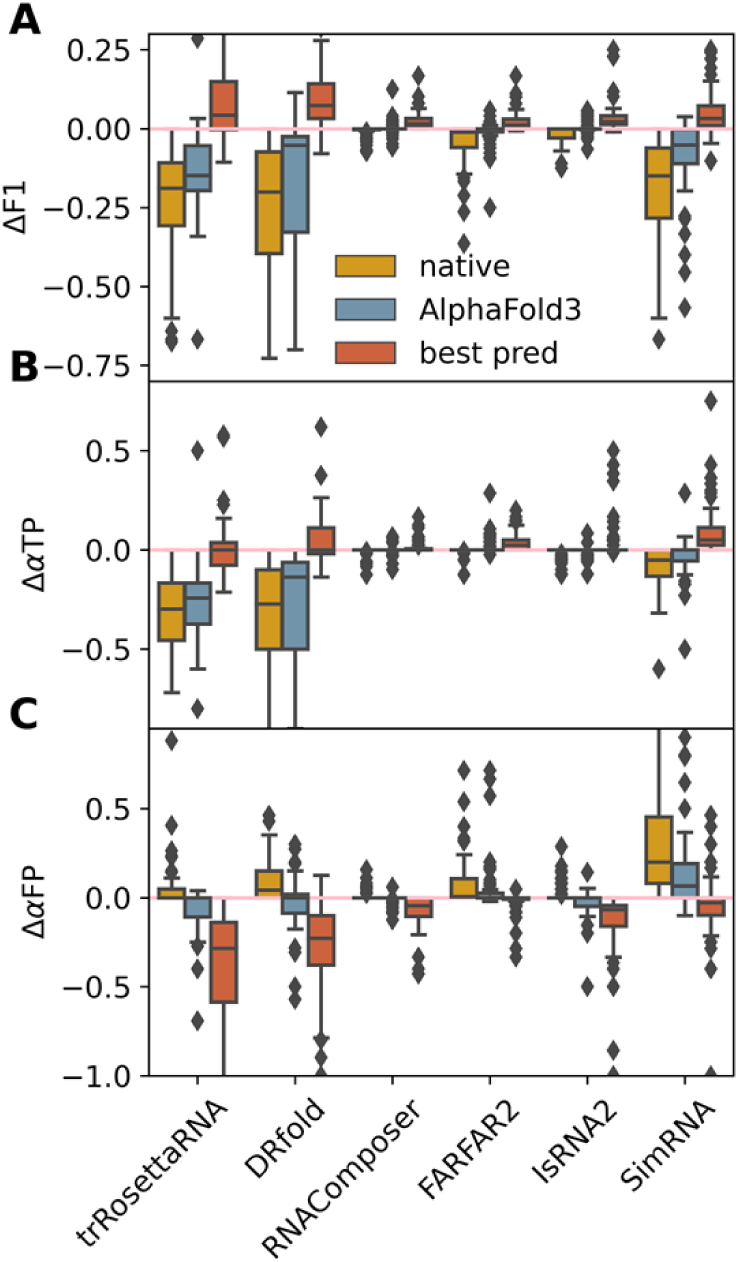
Accuracy of input 2D structures shown distinct effects on the ability of the five tested 3D models to modify base-pairing interactions. Box plots of (A) ΔF1, (B) Δα_*TP*_, and (C) Δα_*FP*_ when using different 2D structures as input for 3D structure predictions. Predictions using the native and AlphaFold3-derived 2D structures as inputs are denoted as “native” and “AlphaFold3”, respectively, while “best pred” represents the best predictions from the optimal combination of 2D and 3D structure prediction models.

## Discussions and conclusions

For large-sized RNAs with complex structural topologies and/or long unpaired loops, accurate 3D structure prediction remains challenging even with the native 2D structure as input. In general, large-sized RNAs often possess complicated interaction networks beyond the standard Watson-Crick base-pairing interactions, making it difficult to accurately describe these interactions *via* scoring/energy functions (known as the scoring problem). In addition, RNA molecules are highly flexible and can fold into a vast number of possible conformations, making it computationally expensive to find an appropriate final structure (known as the sampling problem). As a result, the folding free energy landscape of complex RNA structures is very rugged with multiple local minima, making it challenging for current computational algorithms to identify the global minimum corresponding to the native conformation. Therefore, although we have recently witnessed remarkable progress in deep learning-based structural modeling of biological macromolecules, even the state-of-the-art methods, including the recent Alphafold3 [29], still cannot achieve a level of accuracy in RNA 3D structure modeling comparable to that of protein structure prediction [26,28].

Due to the complexity of RNA folding, many deep learning-based methods and majority of traditional approaches tend to use 2D structures as input to improve the accuracy of 3D structure prediction, where the 2D structures may be generated by existing 2D tools, multiple sequence alignments, or RNA foundation models. Our study found that the cross-talk between the input 2D structure and the resulting 3D structure prediction is prevalent in all tested 3D models. That is, the performance of RNA 3D structure prediction is highly dependent on the accuracy of the input 2D structure, and the 3D models are able to modify the original base-pairing interactions contained in the input 2D structure during the 3D structure modeling process. For instance, 3D structure prediction can improve the interaction network through introducing additional true positive base pairs and eliminating some false positive base pairs in the input 2D structure; and *vice versa*. We also observed a close relationship between the accuracy dependence of 3D structure prediction on input 2D structure and the ability of the 3D models to modify the input base-pairing interactions. For example, the template-based RNAComposer has a weak ability to modify the input base-pairing interactions, so its 3D structure prediction performance is heavily dependent on the accuracy of input 2D structure; while the deep learning-based DRfold has a more significant ability to modify the input base-pairing interactions, especially for input 2D structures of medium and high accuracy, thus its performance dependence on the accuracy of input 2D structure is clearly distinct (see Figs 6 and 7 for details). Particularly, for RNA pseudoknots containing tertiary interactions, DRfold’s predictions using predicted 2D structures as input perform better than those based on the native 2D structure (see Fig 2). Therefore, when using predicted 2D structure of medium accuracy as input, 3D structure predictions by DRfold generally perform better because they usually possess improved base-pairing interactions during structure modeling; when using high-confidence 2D structure (close to the native structure) as input, predictions by RNAComposer and IsRNA2 seem more promising because they preserve the input correct base-pairing interactions well, while DRfold’s predictions usually tend to deteriorate the input interactions.

Regardless, the high-accuracy 2D structure as input is one of the keys to improving the quality of RNA 3D structure prediction. Although there may be an upper limit, when the accuracy of input 2D structure exceeds this limit, it has no effect or even a detrimental impact on the 3D structure prediction. Our study also suggests that the performance of 3D structure prediction is more sensitive to the occurrence of false positive base pairs in input 2D structure than to the true positive base pairs, which was indicated by the larger magnitude of Pearson correlation coefficients for the former in each 3D model (see Figs 4 and S7). In addition, the modification of false positive base-pairing interactions is more likely to occur (see Figs 6 and 7) during the 3D structure modeling process. As in the case of DRfold predictions using the predicted 2D structures as input, the apparently better performance is mainly contributed by the elimination of some false positive base pairs. Therefore, a worthy study direction to further improve the performance of 3D structure prediction is to minimize the occurrence of incorrectly predicted base-pairing interactions (reducing false positives) without compromising the presence of correct interactions (maintaining or slightly increasing true positives). Finally, we anticipate that an iterative procedure integrating 2D and 3D structure predictions may be one of the promising solutions to simultaneously achieve accurate predictions of RNA 2D and 3D structures in the future.

## Methods and Materials

The general process of predicting RNA 3D structure consists of two steps: (i) generating a 2D structure from the given RNA sequence using appropriate 2D structure prediction models, and (ii) predicting the 3D structure using the sequence and the generated 2D structure as input. The RNA 2D and 3D structure prediction tools tested in this work are briefly introduced below.

### 2D structure prediction models

Six popular 2D structure prediction models were used to generate RNA 2D structures from sequence information in this work, including RNAfold [53], RNAStructure [54], CONTRAfold [55], Mfold [56–58], NUPACK [61], and MXfold2 [60]. These six models can be categorized into two groups: thermodynamics-based models and deep learning-based models. Their details are displayed in Table 2. In the following benchmarks, all the prediction packages were downloaded and run locally. For reference, the native 2D structures extracted from experimental structures using DSSR [72] were also prepared as input and denoted as the *native* 2D structure in subsequent tables and figures. In addition, the 2D structures extracted from 3D structures predicted by AlphaFold3 [29] using DSSR were also used as another baseline (denoted as AlphaFold3).

### 3D structure prediction methods

Considering the diversity of prediction categories (deep learning-based, template-based, and *de novo* methods) and their availability at the time of preparing this manuscript, five representative RNA 3D structure prediction methods were benchmarked here, including DRfold [30], RNAComposer [5,46], FARFAR2 [7], IsRNA2 [17], and SimRNA [19]. Here we choose DRfold as the representative deep learning-based method because of its top performance in a recent benchmark of deep-learning methods for RNA 3D structure prediction [50]. Moreover, DRfold does not rely on multiple sequence alignment to prepare input, which could avoid unexpected modifications to the input 2D structure and ensure the purity of our benchmark study. Notably, DRfold utilizes PETfold [73] and RNAfold [53] to predict secondary structure from RNA sequence input during the 3D structure prediction process. In order to enable user-specified RNA 2D structure input, we modified the PETfold_runner and ViennaRNA_runner functions in the scripts/Features.py file to bypass secondary structure prediction and instead directly read this information from input files. A summary of these five methods is shown in Table 1, and their brief features are listed in the supplementary information (SI) Text. For each prediction among the five selected 3D models using different 2D structures as input, the top five predictions were collected by local runs (DRfold, FARFAR2, IsRNA2, and SimRNA) or by the webserver (RNAComposer). The best prediction among the five candidates was used for subsequent studies. Additionally, we also considered the AlphaFold3 [29] and trRosettaRNA [33] predictions as a reference. The webserver at https://alphafoldserver.com/ with default settings was used for AlphaFold3, while the trRosettaRNA run locally. Their brief features are also listed in the supplementary information (SI) Text.

### Test datasets

To comprehensively evaluate the performance of five representative RNA 3D structure prediction methods, we constructed three test datasets, including “Custom”, “RNA Puzzles”, and “CASP RNA”. From the protein data bank (PDB) (as of December 2024), a Custom dataset of 31 RNA molecules was prepared by clustering sequences with Cd-hit [74] at an 80% sequence similarity threshold and excluding those appearing in the DRfold training set. These 31 RNAs have varied lengths and structural topologies; for more details, see Table S1 in the SI. The RNA Puzzles dataset contains 18 RNAs collected from the real challenging RNA-Puzzles [38–42], excluding cases that appeared in the DRfold training set; see Table S2 for more details. The CASP RNA dataset covers 14 RNA targets used in the recent CASP 15 and 16 competitions; see details listed in Table S3. Moreover, we merged these three test datasets (Custom, RNA Puzzles, and CASP RNA) to form a Combined dataset (63 RNAs). For in-depth analysis, we also reclassified the Combined dataset into three categories according to the structural topology, namely stem-loop (22 RNAs), multi-way junction (16 RNAs), and pseudoknot (25 RNAs). As shown in Fig S1, based on the native 2D structure, a nearly linear relationship between the number of nucleotides (*N*_*nt*_) and the number of base pairs (*N*_*pair*_) was observed in the Combined dataset, indicating that almost all tested RNAs formed substantial 2D structures. In terms of structural topology, the proportions of stem-loop, multi-way junction, and pseudoknot (containing tertiary interactions) RNAs are 34.9%, 25.4%, and 39.7%, respectively, declaring the richness and balance of structural types in the test datasets.

### Evaluation metrics

Same as in previous studies [60,62], the F1-score was used to evaluate the accuracy of RNA 2D structure prediction, with values between [0, 1] and F1-score = 1 indicating that the predicted 2D structure perfectly matches the native one. To comprehensively assess the accuracy of RNA 3D structure predictions, we considered multiple metrics including Root-Mean-Square-Deviation (RMSD) [75], Interaction Network Fidelity of all base-base interactions (INF_ALL) [76], Local Distance Difference Test (lDDT) [77], and TMscore [78]. See SI Text for their detailed definitions. Briefly, the values of INF_ALL, TMscore, and lDDT range from 0 to 1, where value of one indicates an ideal prediction. To further investigate the impact of the input 2D structure (base pairs) on the accuracy of 3D structure prediction, three additional metrics were introduced, namely the proportion of true positive (α_*TP*_), false positive (α_*FP*_), and false negative (α_*FN*_) predictions, defined as

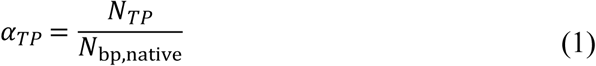

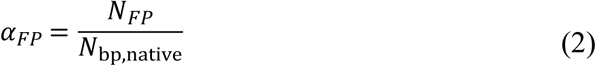

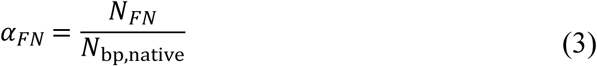

where *N*_*TP*_, *N*_*FP*_, and *N*_*FN*_ represent the number of true positive, false positive, and false negative predicted base pairs, respectively. *N*_bp,native_ denotes the number of base pairs contained in the native 2D structure. The proportions α_*TP*_ and α_*FN*_ take values in the range [0, 1], while α_*FP*_ may exceed than 1.

## Data availability

Details of the test datasets are included in Supporting Information Tables S1-S3. Prediction results are available at https://github.com/DongZhangRNA/2D-and-3D-benchmark for community use.

## Supporting information

**S1 File.** Supplementary text. (DOCX)

**S1 Fig** (A) Pair plots of the number of the nucleotides (*N*_*nt*_) and the number of the base pairs (*N*_*pair*_) in the Combined dataset. (B) Pie chart for RNA structure topologies in the Combined dataset: stem-loop, multi-way junction, and pseudoknot (containing tertiary interactions). (C) Distributions of the proportions of paired nucleotides (^2*N*^*pair* / ^*N*^*nt*) in different structure topologies.

(TIFF)

**S2 Fig** Box plots of (A) RMSD, (B) INF_ALL, and (C) TMscore for different combinations of 2D and 3D structure predictions models in the Combined dataset. Individual 2D structure predicted by six popular 2D tools, the 2D structure derived from 3D structure predicted by AlphaFold3, or the native 2D structure was used as input for five selected 3D models to predict RNA 3D structure. From left to right: DRfold, RNAComposer, FARFAR2, IsRNA2, and SimRNA.

(TIFF)

**S3 Fig** Box plots of (A) RMSD, (B) INF_ALL, and (C) TMscore for predictions from five tested 3D models using different 2D structures as input: stem-loops (first column), multi-way junctions (second column), and pseudoknots (third column). Predictions using native 2D structures and 2D structures derived from 3D structures predicted by AlphaFold3 as inputs are denoted as “native” and “AlphaFold3”, respectively, while “best pred” represents the best predictions from the optimal combination of 2D and 3D structure prediction models.

(TIFF)

**S4 Fig** The proportion of different topological categories as functions of the F1-score values of input 2D structures. The F1-score values were grouped by a bin size of 0.1. Data represent the mean value of a particular bin.

(TIFF)

**S5 Fig** Detailed relationship between the (A-C) RMSD and (D-F) INF_ALL values of RNA 3D structures predicted by the five selected models and the F1-score values of input 2D structures on different test datasets: (A, D) Custom dataset, (B, E) RNA Puzzles, and (C, F) CASP RNA. The F1-score values were grouped by a bin size of 0.1. Symbols represent the mean metrics of a particular bin and shaded area indicates the 95% confidence interval. The analysis considered all 2D structures predicted by the selected six 2D tools, AlphaFold3-derived 2D structures, and the native 2D structures.

(TIFF)

**S6 Fig** Accurate 2D structures as input can largely improve the performance of RNA 3D structure prediction. Box plots of (A) RMSD, (B) INF_ALL, and (C) TMscore values of RNA 3D structure predictions based on different accuracy levels of input 2D structures: low accuracy (F1-score < 0.5), medium accuracy (0.5 ≤ F1-score < 0.9), and high accuracy (F1-score ≥ 0.9).

(TIFF)

**S7 Fig** Scatter plots shown the relationships between the (A) proportion of true positive

(α_*TP*_) base pairs and (B) proportion of false positive (α_*FP*_) base pairs in the input 2D structure and the RMSD values of 3D structures predicted by different 3D models. The line in each plane represents the linear fitting result, and the corresponding Pearson correlation coefficient (*R*) and p-value (*p*) are also presented. From left to right: DRfold, RNAComposer, FARFAR2, IsRNA2, and SimRNA.

(TIFF)

**S8 Fig** (A) Native 2D structure of the crRNA in the CRISPR-Cas Phi system (PDB id: 7M5O [79]). Using the native 2D structure as input, the 3D structures predicted by different 3D models: (B) DRfold, (C) RNAComposer, (D) FARFAR2, (E) IsRNA2, and (F) SimRNA. INF_ALL and RMSD values between the predicted (in red) and native (in blue) structures for each model are also shown.

(TIFF)

**S9 Fig** Histogram of the fraction of F1-score after 3D structure prediction (F1_3*D*_) greater than, equal to, and lower than the F1-score of the input 2D structure (F1_2*D*_) for each tested 3D model. Here all the combination of selected six 2D and five 3D methods were considered.

(TIFF)

**S1 Table.** List of RNAs in the Custom dataset. (XLSX)

**S2 Table.** List of RNAs in the RNA Puzzles dataset. (XLSX)

**S3 Table.** List of RNAs in the CASP RNA dataset. (XLSX)

**S4 Table.** The full table of RMSD results. (XLSX)

## Author Contributions

**Conceptualization:** Dong Zhang.

**Data curation:** Deyin Wang, Yangwei Jiang.

**Formal analysis:** Deyin Wang, Yangwei Jiang, Dong Zhang, Ruhong Zhou, Linli He, Linxi Zhang.

**Funding acquisition:** Dong Zhang, Linli He, Linxi Zhang.

**Investigation:** Deyin Wang, Yangwei Jiang.

**Methodology:** Deyin Wang, Yangwei Jiang, Dong Zhang, Linli He, Linxi Zhang, Ruhong Zhou.

**Project administration:** Dong Zhang, Linli He, Linxi Zhang.

**Resources:** Dong Zhang, Linli He, Linxi Zhang, Ruhong Zhou.

**Supervision:** Dong Zhang, Linli He, Linxi Zhang.

**Validation:** Deyin Wang, Yangwei Jiang, Dong Zhang, Linli He, Linxi Zhang, Ruhong Zhou.

**Visualization:** Deyin Wang, Yangwei Jiang.

**Writing** – **original draft:** Deyin Wang, Yangwei Jiang.

**Writing** – **review & editing:** Deyin Wang, Yangwei Jiang, Dong Zhang, Linli He, Linxi Zhang, Ruhong Zhou.

